# Functional characterization of helminth-associated Clostridiales reveals covariates of Treg differentiation

**DOI:** 10.1101/2023.06.05.543751

**Authors:** Shushan Sargsian, Alannah Lejeune, Defne Ercelen, Wen-Bing Jin, Alan Varghese, P’ng Loke, Yvonne A. L. Lim, Chun-Jun Guo, Ken Cadwell

## Abstract

Parasitic helminths influence the composition of the gut microbiome. However, the microbiomes of individuals living in helminth-endemic regions are understudied. The Orang Asli, an indigenous population in Malaysia with high burdens of the helminth *Trichuris trichiura*, displayed microbiotas enriched in Clostridiales, an order of spore-forming obligate anaerobes previously shown to have immunogenic properties. We previously isolated novel Clostridiales that were enriched in these individuals and found that a subset promoted the *Trichuris* life cycle. Here, we further characterized the functional properties of these bacteria. Enzymatic and metabolomic profiling revealed a range of activities associated with metabolism and host response. Consistent with this finding, monocolonization of mice with individual isolates identified bacteria that were potent inducers of regulatory T cell (Treg) differentiation in the colon. Comparisons between variables revealed by these studies identified enzymatic properties correlated with Treg induction and *Trichuris* egg hatching. These results provide functional insights into the microbiotas of an understudied population.

## Introduction

The gut microbiota and its enzymatic byproducts impact numerous aspects of host physiology including metabolism and immunity [1–9]. The composition of the microbiota has been associated with numerous disease conditions and is heavily influenced by geography and lifestyle [10–18]. However, microbiome research is dominated by studies examining individuals in highly developed countries [19, 20]. Environmental variables absent in these populations contribute to differences in their microbiome compared with individuals residing in other regions. For instance, parasitic worms known as helminths cohabitate the gastrointestinal tract alongside bacteria and influence the diversity and composition of the microbiota [21–25]. Helminths colonize 1.5 billion people, or roughly 24% of the world population, with the highest prevalence reported in tropical and subtropical areas in Africa, Asia, and South America. [26–28]. Examining the properties of intestinal bacteria enriched in individuals from helminth-endemic regions may broaden our understanding of microbiota function.

Helminth infection can cause disease involving malnutrition, impaired growth, dysentery, intestinal obstruction, and anemia [26, 27]. The incidence of helminth infections is also negatively linked to the incidence of immune-mediated disorders on a global scale [29, 30]. The immune response generated against helminths includes the differentiation and expansion of regulatory T cells (Tregs), which suppress the development of autoimmune and inflammatory diseases [31–36]. Immune modulation by helminths can occur either by direct effects on the host immune system or indirectly through the microbiota [37–41]. For example, we and others have shown that expansion of the Clostridiales order during helminth colonization ameliorates disease in mouse models of inflammatory bowel disease and allergic asthma [41, 42]. We also found that Clostridiales were enriched in the gut microbiome of the Orang Asli, an indigenous population in rural Malaysia. Clostridiales were enriched in individuals with high burdens of the whipworm *Trichuris trichiura*, and deworming medication reduced the relative abundance of Clostridiales [42].

Clostridiales and the Clostridia class to which they belong are Gram-positive, spore-forming anaerobes that influence host physiology, such as through promoting differentiation of Tregs [5, 43, 44]. Based on the above association with helminths, we recently used a chloroform-based enrichment protocol to isolate and sequence the genomes of spore-forming bacteria from the feces of helminth-colonized Orang Asli [45]. This approach identified 13 Clostridiales species, most of which represented poorly characterized taxa. Metagenomics analysis of a large number of Malaysians confirmed the association of these Clostridiales members with the Orang Asli population and identified a specific association between the *Peptostreptococcaceae* family and helminth colonization. Also, we found that *Peptostreptococcaceae* isolates were strong inducers of *Trichuris* egg hatching, suggesting that Clostridiales enriched during helminth colonization contain properties that support the helminth life cycle. However, the functional properties of these bacteria remain unknown. Here, we examined the enzymatic and metabolomic activity of isolated bacteria and their ability to induce Treg differentiation and identified specific enzymatic properties that correlated with Treg induction and *Trichuris* egg hatching.

## Results

### Study design

We previously isolated 14 distinct bacterial taxa from *T. trichiura*-colonized Orang Asli (OA1-14) and assigned genus-species designation based on whole genome sequencing (Figure 1) [45]. OA isolates represented six Clostridiales families: *Erysipelotrichaceae*, *Coprobacillaceae*, *Clostridiaceae*, *Peptostreptococcaceae*, *Oscillospiraceae*, and *Lachnospiraceae*. One of the *Lachnospiraceae* members, OA3, was a new species in the *Ruminococcus* genus based on sequence identity and was designated *Ruminococcus pangsunibacterium*. OA12 was identified as *Enterococcus hirae*. Although not a member of the Clostridiales order, the resistance of *Enterococci* to chemical treatment likely explains how this bacterium was retained after fractionation [46, 47]. Similar to the other OA isolates, OA12 was enriched in Malaysian microbiomes [45] and was therefore included in subsequent experiments. OA isolates were analyzed for *in vitro* enzymatic and metabolomic activity, and then screened for their capacity to promote Treg differentiation *in vivo*. Finally, we performed correlation analyses to look for relationships between all measured variables to identify bacterial properties related to Treg differentiation or *Trichuris* hatching.

**Figure 1.**
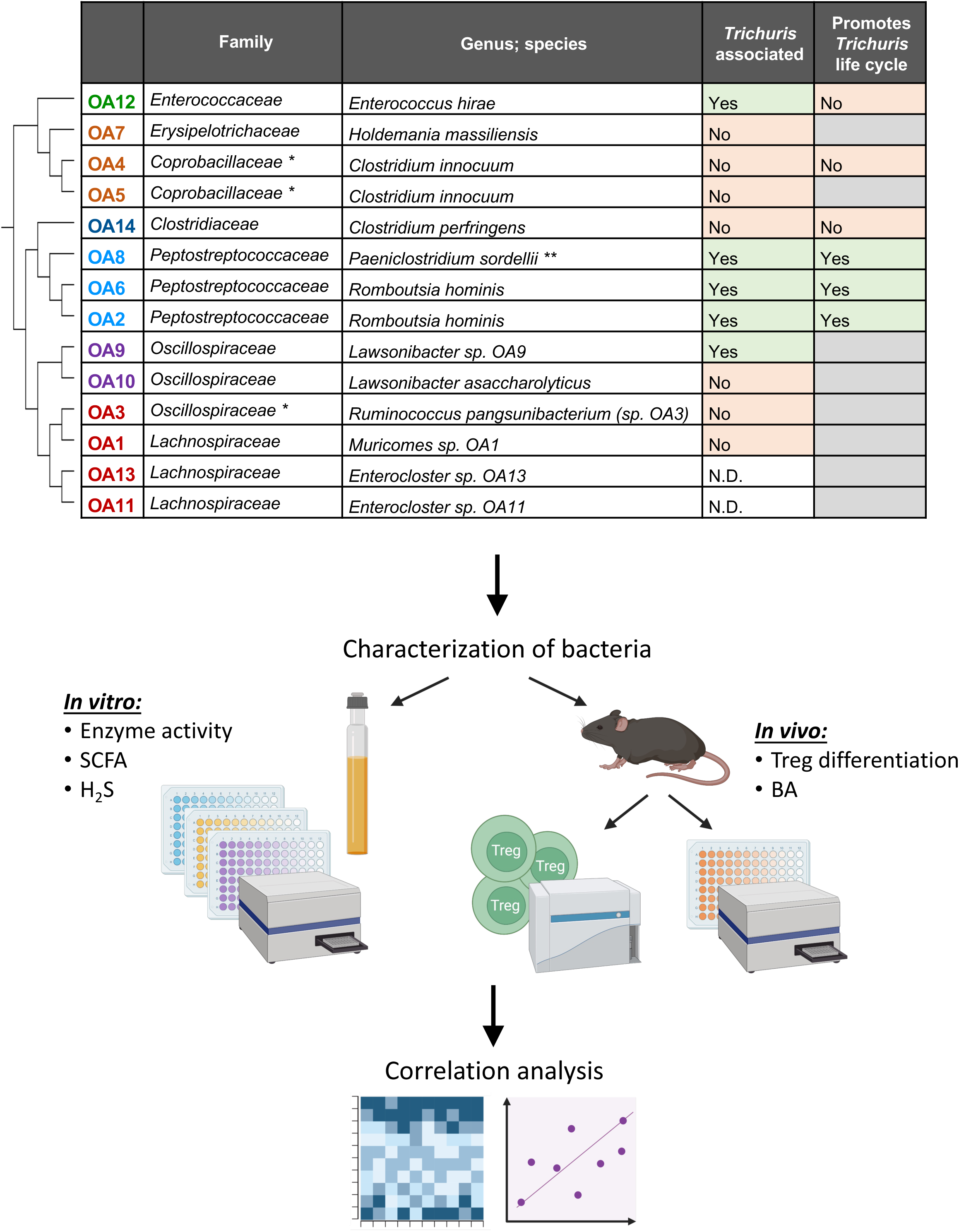
Study design for characterization of the OA isolates. List of OA isolates with updated taxonomic classifications (top) and schematic summarizing the study design (bottom). The phylogenetic tree to the left of the table depicts the relationships between OA isolates. The column labeled “*Trichuris* associated” refers to whether the abundance of the isolate decreased in microbiomes after deworming treatment, with ‘N.D.’ denoting isolates that were not detected [45]. The column labeled “Promotes *Trichuris* life cycle” refers to whether the isolate promotes *Trichuris muris* egg hatching, with gray cells denoting isolates that were not tested [45]. * OA4 and OA5 were classified as members of the *Erysipelotrichaceae* family and OA3 was classified as a member of the *Lachnospiraceae* family at the time of our previous publication [45]. ** OA8 was classified as *Paraclostridium sordellii* at the time of our previous publication [45]. SCFA, short chain fatty acids; H_2_S, hydrogen sulfide; BA, bile acids.

### Enzymatic profile of the OA isolates

OA isolates were functionally profiled for their ability to perform 57 enzymatic reactions that represent common bacterial activities including carbohydrate metabolism. Taxonomically related isolates were more likely to share activities (Figure 2a-c). For instance, *Peptostreptococcaceae* isolates (OA2, 6 and 8) and *Clostridium perfringens*-OA15 displayed high levels of acid phosphatase, esculin hydrolysis, Naphthol-AS-BI-phosphohydrolase, and glutamic acid decarboxylase activities (Figure 2a) and the ability to metabolize D-glucose (Figure 2c). The *Peptostreptococcaceae* isolates, in addition to *E. hirae*-OA12, also had the highest arylamidase activities, both in terms of the degree of activity and the different amino acid substrates (Figure 2b). As expected, the two *Clostridium innocuum* isolates OA4 and OA5 displayed similar profiles for the majority of the 57 parameters, although there were a few enzymatic reactions in which they diverged. For instance, *C. innocuum*-OA4 but not *C. innocuum*-OA5 showed mannose and raffinose fermentation activities. *E. hirae*-OA12 and *Enterocloster* species OA11 and OA13 displayed the broadest capacities to metabolize carbohydrates (Figure 2c). Overall, these data indicate that OA isolates possess a broad range of metabolic activities, consistent with their taxonomic diversity.

**Figure 2.**
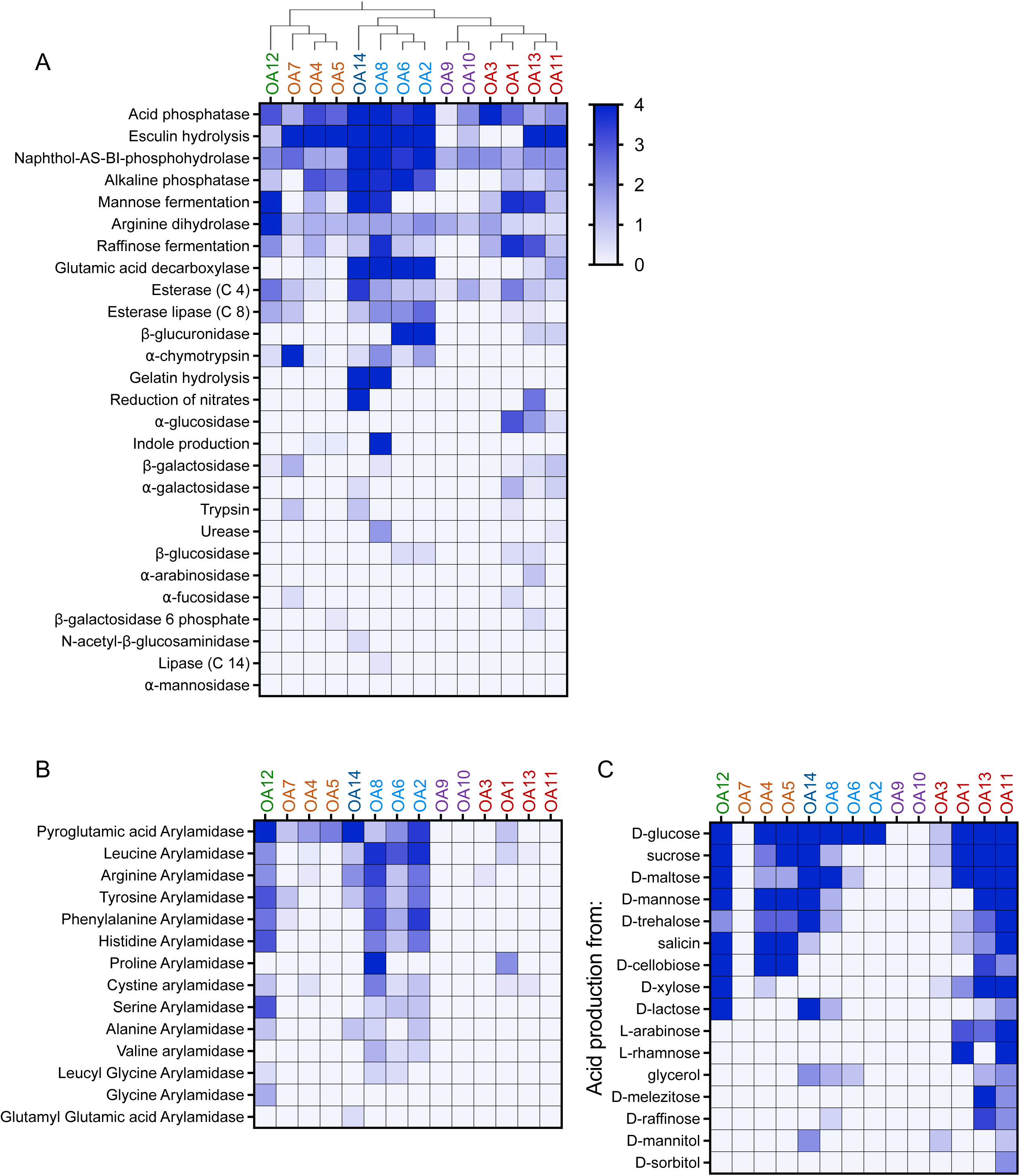
Enzymatic profile of the OA isolates. Enzyme activity (A), arylamidase activity (B), and acid production from various carbohydrates (C) by the OA isolates were measured using API colorimetric kits for microbial identification. Each reaction was run in at least three independent trials for each isolate, and the intensity of the reaction was assessed visually on a scale of 0 (no reaction) to 4 (strong or complete reaction). The phylogenetic tree at the top of (A) depicts the relationships between OA isolates.

### OA isolates produce short chain fatty acids and hydrogen sulfide

Given the link between helminth-associated microbiota and anti-inflammatory responses, we measured the production of short chain fatty acids (SCFAs) and hydrogen sulfide (H_2_S) by the OA isolates. Bacteria ferment soluble fibers to produce short chain fatty acids (SCFAs) which include acetate, propionate, and butyrate [48]. SCFAs have been shown to regulate a variety of immune cell types [9]. For example, they can induce Treg expansion in the gut to regulate intestinal inflammation [49–51]. Although levels varied, we detected acetate in the culture supernatant for all 14 OA isolates. *C. innocuum*-OA4 and -OA5 and *C. perfringens*-OA14 produced butyrate at levels similar to or greater than acetate. Propionate was not detected or negligible except for *Paraclostridium sordellii*-OA8, the two *Romboutsia hominis*-OA2 and -OA6, and *Enterocloster sp*. OA11 (Figure 3a). We validated this finding by testing cecal samples from mice monocolonized with *R. hominis*-OA2 and included germ-free (GF) mice as a negative control and mice colonized with a previously described synthetic minimal flora (MF) consisting of 15 bacteria [52, 53] as a positive control. As expected, we detected minimal levels of SCFAs in GF mice compared with MF mice in which all three were detected. Mice monocolonized with OA2 displayed the proportion of the three SCFAs predicted by the *in vitro* analysis (Figure 3b).

**Figure 3.**
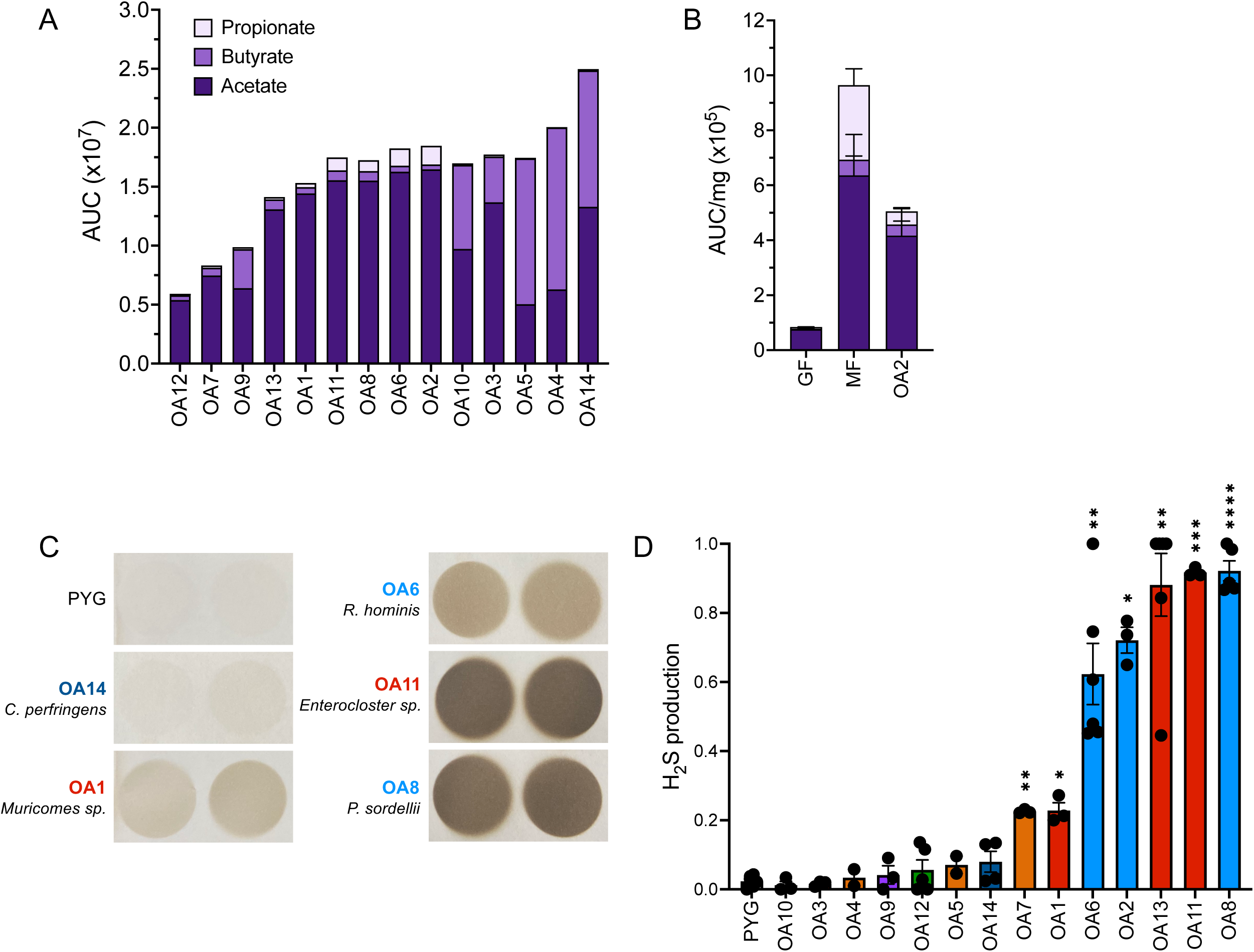
OA isolates produce short chain fatty acids and hydrogen sulfide. (A) Short chain fatty acid (SCFA) production in bacterial culture supernatants measured by mass spectrometry. AUC, Area under curve. (B) SCFA concentration per mg of cecal contents from germ-free (GF), minimal flora (MF) or *R. hominis*-OA2-monocolonized (OA2) mice, measured by mass spectrometry. AUC, Area under curve. (C) Representative images of lead acetate paper used to detect hydrogen sulfide (H_2_S) production by the OA isolates *in vitro*. Darker color indicates more H_2_S produced by bacterial cultures. (D) Quantification of H_2_S production by each OA isolate, normalized to the amount of H_2_S production by media alone (‘PYG’) within each experiment. Kruskal-Wallis with Dunn’s multiple comparisons test was used to compare between each group and PYG. *p < 0.05, **p < 0.01, ***p < 0.001, ****p < 0.0001.

Some bacteria can produce hydrogen sulfide (H_2_S), which suppresses inflammation in multiple disease models and is associated with Treg differentiation [54–59]. We found that *Peptostreptococcaceae* isolates OA2, 6 and 8, as well as *Enterocloster* isolates OA11 and OA13, were potent producers of H_2_S, suggesting another possible mechanism by which these bacteria could have immunomodulatory interactions with the host (Figure 3c-d).

### Several OA isolates induce regulatory T cells in the gut

Peripherally induced Tregs (iTregs) in the gut that develop in response to microbial antigens and metabolites are distinguished by expression of the transcription factor *forkhead box P3* (*Foxp3*) and *RAR-related orphan receptor γt* (*Rorγt*). These Foxp3^+^ Rorγt^+^ Tregs contribute to immune tolerance of the microbiota and suppress colitis [60–64]. iTregs can be further distinguished from thymus-derived or natural Tregs (nTregs) reactive to self-antigens by the absence of the transcription factor Helios (Ikzf2) [65]. GF mice possess low numbers of iTregs, and thus, the induction of iTregs in GF mice in response to colonization by bacteria serves as a sensitive assay to measure immunomodulatory potential of commensal species [61, 66–68].

We analyzed iTregs in the colonic lamina propria of GF mice by flow cytometry three to four weeks post oral inoculation with individual OA isolates (Figure 4a), except *Lawsonibacter* sp. OA9 which failed to colonize GF mice in the absence of other bacteria. In addition to conventional specific pathogen free (SPF) mice and untreated GF mice that served as benchmarks, we included several controls to aid in interpretation of the quantitative data. As a positive control, we colonized mice with a mixture of 17 Clostridia isolates (KH mix) previously shown to induce Tregs in GF mice [5]. As negative controls, we used mice monocolonized with segmented filamentous bacteria (SFB), a bacterium that induces Th17 cells (Rorγt^+^ Foxp3^-^ CD4^+^ T cells) without inducing Tregs [69], and mice monocolonized with one of the KH mix strains, *Clostridium aldenense* (KH28), which is insufficient to induce Tregs to levels achieved with the entire KH mix consortium [5]. We also quantified Tregs in MF mice described above (mice raised in a gnotobiotic isolator containing MF) and GF mice gavaged with stool from MF mice (MF stool). Stable colonization was confirmed for all conditions based on detection of bacteria in stool (Figure S1A).

**Figure 4.**
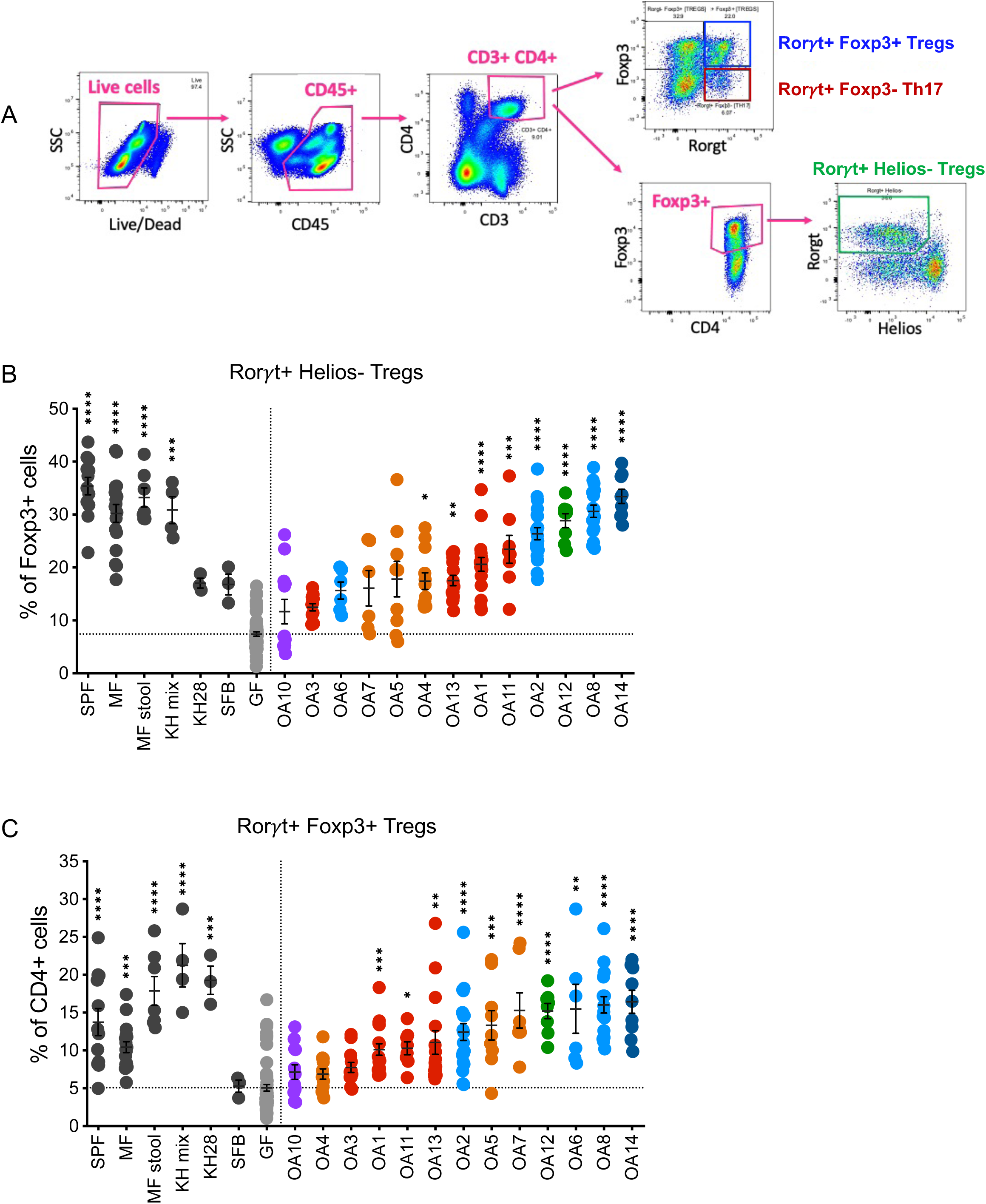
OA isolates induce iTregs in the colon. (A) Representative flow cytometry gating strategy for Rorγt^+^ Helios^-^ iTregs out of Foxp3^+^ cells, Foxp3^+^ Rorγt^+^ iTregs out of CD4^+^ cells, and Rorγt^+^ Foxp3^-^ Th17 cells out of CD4^+^ cells from colon lamina propria cells. (B) Frequencies of Rorγt^+^ Helios^-^ cells within Foxp3^+^ Tregs. (C) Frequencies of Rorγt^+^ Foxp3^+^ Tregs out of CD4^+^ T-cells. SPF, Specific pathogen free; MF, Minimal flora mice; MF stool, GF mice gavaged with stool from Minimal flora mice; KH mix, Mice gavaged with consortium of 17 Clostridia strains from Atarashi et al., 2013 [5]; KH 28, *Clostridium aldenense* from the KH mix; SFB, Segmented filamentous bacteria; GF, Germ-free. Each dot corresponds to one mouse. Each group was tested in at least 2 independent experiments consisting of 2-5 mice. Kruskal-Wallis with Dunn’s multiple comparisons test was used to compare between each group and GF. *p < 0.05, **p < 0.01, ***p < 0.001, ****p < 0.0001.

As expected, GF mice had lower frequencies of Rorγt^+^ Helios^-^ Tregs out of all Foxp3^+^ cells and lower Foxp3^+^ Rorγt^+^ Tregs out of CD4^+^ cells compared to SPF, MF, MF stool and KH mix groups (Figure 4b-c). We observed a spectrum of Treg induction by the OA isolates. The proportion of Foxp3^+^ Rorγt^+^ Tregs and Foxp3^+^ Rorγt^+^ Helios^-^ Tregs were restored to similar levels as SPF or MF mice in mice monocolonized with several of the OA isolates including *P. sordellii*-OA8, *E. hirae*-OA12, and *C. perfringens*-OA14. In contrast, several OA isolates such as *R. pangsunibacterium*-OA3 and *Lawsonibacter asaccharolyticus*-OA10 were similar to SFB and failed to induce significant increases in these Treg populations. Mice monocolonized with KH28 increased the frequency of Foxp3^+^ Rorγt^+^ Tregs out of CD4^+^ cells but not Rorγt^+^ Helios^-^ Tregs out of Foxp3^+^ cells (Figure 4b-c). In addition to changes in the proportion of these populations, we quantified the absolute number of Tregs per colon. This analysis largely confirmed the above findings by showing that OA isolates that increased the relative frequency of Foxp3^+^ Rorγt^+^ Tregs and Foxp3^+^ Rorγt^+^ Helios^-^ Tregs also increased the total number of these cells (Figure S1b-c).

### Treg induction by OA isolates correlates with Th17 but not colonization levels

Given their role in mucosal immunity [70, 71], we examined whether OA isolates affect the proportion of Th17 cells (Rorγt^+^ Foxp3^-^ CD4+ T cells) (Figure 4a) in the gut. Similar to Tregs, we found that monocolonization with OA isolates led to a range of Th17 cell levels in the colon, although none reached the extraordinarily high levels detected in mice monocolonized with SFB (Figure 5a, S1d). Excluding SFB, we found that the proportion of Th17 cells in the colon roughly correlated with the proportion of Tregs (Figure 5b). There were notable exceptions, such as OA6, which was a strong Treg inducer but did not increase the frequency of Th17 cells above levels seen in GF mice.

**Figure 5.**
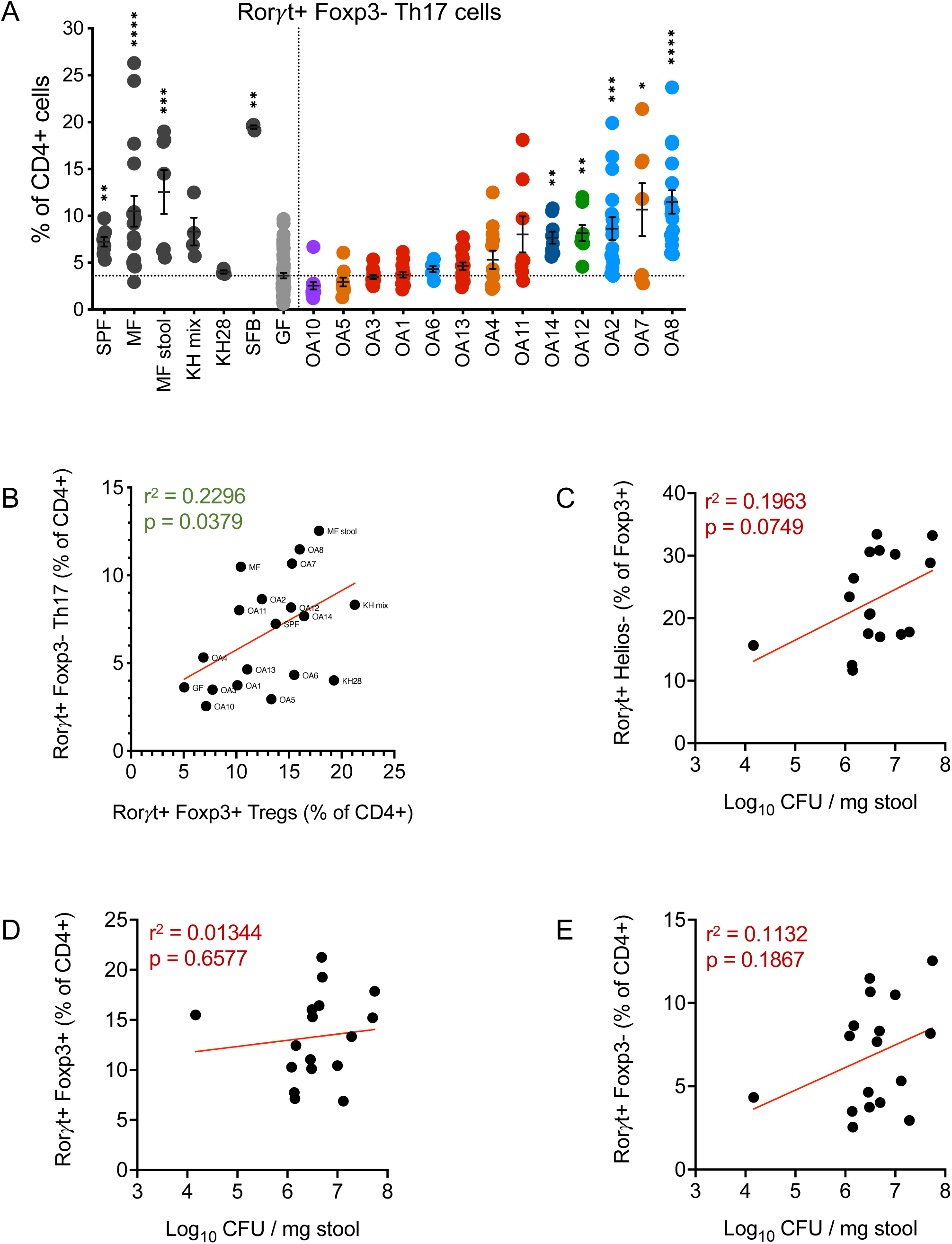
Associations between OA isolate colonization, Tregs, and Th17 cell levels. (A) Frequencies of Rorγt^+^ Foxp3^-^ Th17 cells out of CD4^+^ T-cells. (B) Spearman correlation between the frequency of Foxp3^+^ Rorγt^+^ iTregs out of CD4^+^ cells and Rorγt^+^ Foxp3^-^ Th17 cells out of CD4^+^ cells induced by each bacterium or control group. (C-E) Spearman correlation between the bacterial burden in each mouse and the proportions of Rorγt^+^ Helios^-^ iTregs out of Foxp3^+^ cells (C), Foxp3^+^ Rorγt^+^ iTregs out of CD4^+^ cells (D), or Rorγt^+^ Foxp3^-^ Th17 cells out of CD4^+^ cells (E). In A: SPF, Specific pathogen free; MF, Minimal flora mice; MF stool, GF mice gavaged with stool from Minimal flora mice; KH mix, Mice gavaged with consortium of 17 Clostridia strains from Atarashi et al., 2013 [5]; SFB, Segmented filamentous bacteria; GF, Germ-free. Each dot corresponds to one mouse. Each group was tested in at least 2 independent experiments consisting of 2-5 mice. Kruskal-Wallis with Dunn’s multiple comparisons test was used to compare between each group and GF. *p < 0.05, **p < 0.01, ***p < 0.001, ****p < 0.0001. In B-E: r^2^ and p values are depicted on each plot. Each dot corresponds to the average of at least 2 independent repeats with 2-5 mice within each experiment and the red line depicts the linear regression slope.

These observations raised the possibility that T cell differentiation reflected the degree of colonization by bacteria. However, there were no significant correlations between the bacterial burden and the proportions of Tregs or Th17 cells (Figure 5c-e), suggesting that the level of T cell differentiation is a specific property of each bacterium. Interestingly, we found that colonization by some OA isolates increased total bile acids (BAs) detected in the stool compared with GF mice (Figure S2). Transformation of BAs by gut bacteria is associated with differentiation of Tregs and Th17 cells [72–75]. However, we did not observe a correlation between total BAs and these T cell populations.

### Correlations between properties of OA isolates

To identify relationships between the properties measured for each OA isolate, we performed a spearman correlation analysis between all measured variables (Figure S3). This analysis identified several enzymatic activities that were correlated with T cell differentiation (Figure 6a). Induction of Tregs was positively correlated with alanine arylamidase and tyrosine arylamidase activity by bacteria (Figure 6a-e). Th17 cell induction was also positively correlated with tyrosine arylamidase activity, and additionally associated with ⍺-chymotrypsin activity (Figure 6a, f-i). Alanine arylamidase and tyrosine arylamidase have not previously been linked to Treg or Th17 cell biology. ⍺-Chymotrypsin is a member of the serine protease family and has been shown to have antibacterial properties [76], and is also made by pathogenic bacteria in the *Vibrio* genus [77]. Whether this correlation reflects a role of ⍺-chymotrypsin in Th17 cell induction or merely reflects the pro-inflammatory capacity of bacteria that produce this protease and induce Th17 cells through other mechanisms remains to be determined.

**Figure 6.**
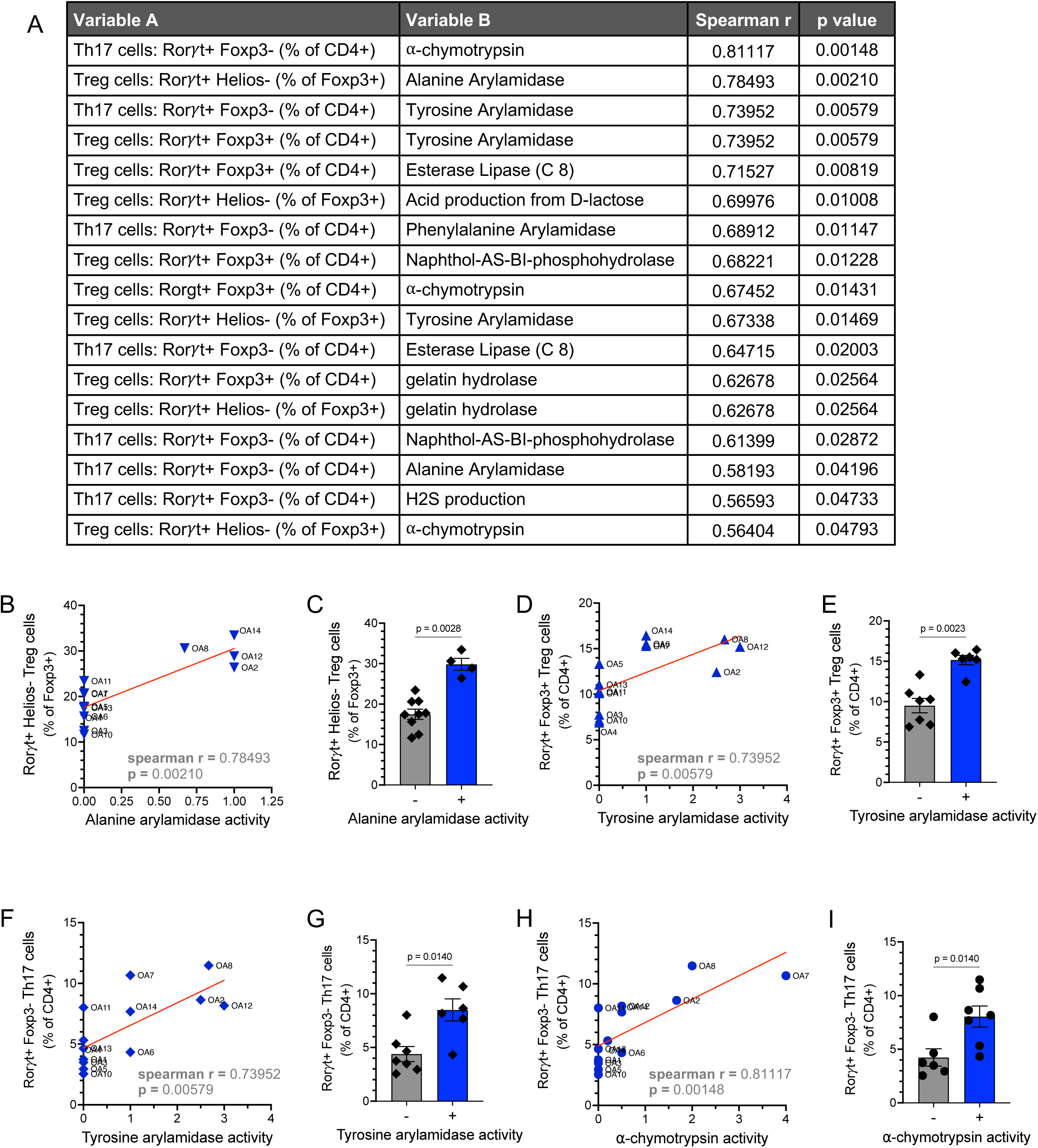
Treg and Th17 cell induction by the OA isolates is correlated with enzymatic activities by bacteria. (A) Spearman r and p values corresponding to significant correlations between the frequency of induced Treg and Th17 cells and other measured variables of the OA isolates. (B) Graph displaying relationship between alanine arylamidase activity and induction of Rorγt^+^ Helios^-^ Foxp3^+^ iTregs for the OA isolates. (C) Quantification of Rorγt^+^ Helios^-^ Foxp3^+^ iTregs comparing OA isolates with or without alanine arylamidase activity. (D) Graph displaying relationship between tyrosine arylamidase activity and induction of Rorγt^+^ Foxp3^+^ Tregs for the OA isolates. (E) Quantification of Rorγt^+^ Foxp3^+^ Tregs comparing OA isolates with or without tyrosine arylamidase activity. (F) Graph displaying relationship between tyrosine arylamidase activity and induction of Rorγt^+^ Foxp3^-^ Th17 cells for the OA isolates. (G) Quantification of Rorγt^+^ Foxp3^-^ Th17 cells comparing OA isolates with or without tyrosine arylamidase activity. (H) Graph displaying relationship between ⍺-chymotrypsin activity and induction of Rorγt^+^ Foxp3^-^ Th17 cells by the OA isolates. (I) Quantification of Rorγt^+^ Foxp3^-^ Th17 cells comparing OA isolates with or without ⍺-chymotrypsin activity. In B, D, F, H: Spearman r and p values are depicted on each plot in. Each dot corresponds to the average value per OA isolate and the red line depicts the linear regression slope. Bar graphs show mean +/- SEM. In C, E, G, I: Each dot corresponds to the average value per OA isolate. Mann-Whitney test with p values depicted on the bar graphs.

To quantify how bacterial properties explain the variance in lymphocyte differentiation we performed a distance-based redundancy analysis (dbRDA) using the characteristics found to be most significantly related with Treg or Th17 induction according to the Spearman correlation analysis and LASSO regression (Figure S4a-b, Table S1). The analysis revealed that these variables together contribute to the variance of responses in Treg and Th17 induction, with the variation in alanine arylamidase production accounting for 41.95% of the variance, and all other variables accounting for less than 20% of the variance each (Figure S4c).

We also incorporated data from our previous study in which we quantified the ability of the OA isolates to mediate hatching of eggs from *T. trichiura* and the mouse parasite *Trichuris muris* [45]. Hatching of *T. trichiura* but not *T. muris* was positively correlated with H_2_S production and negatively correlated with salicin acidification by the OA isolates (Figure 7a-c). H_2_S, in addition to being involved in a wide variety of physiological processes, has been shown to promote hatching of eggs from *Ascaris suum*, a helminth that infects pigs [78]. Salicin utilization by bacteria was previously shown to be toxic to nematodes in soil [79], although these experiments were not done with *Trichuris* species. Our finding that these bacterial properties are significantly related to egg hatching of only the human whipworm could provide insight on mechanisms that may differentiate dependencies of *T. trichiura* and *T. muris* on bacterial factors for hatching. Valine arylamidase activity was significantly correlated with hatching of both *T. muris* and *T. trichiura* eggs (Figure 7a, d-e). Valine arylamidases were among the proteases found to be part of the excretory/secretory products of *Contracaecum rudolphii*, a nematode that infects mammals and birds [80]. This observation suggests a role for this protease in *Trichuris* egg hatching.

**Figure 7.**
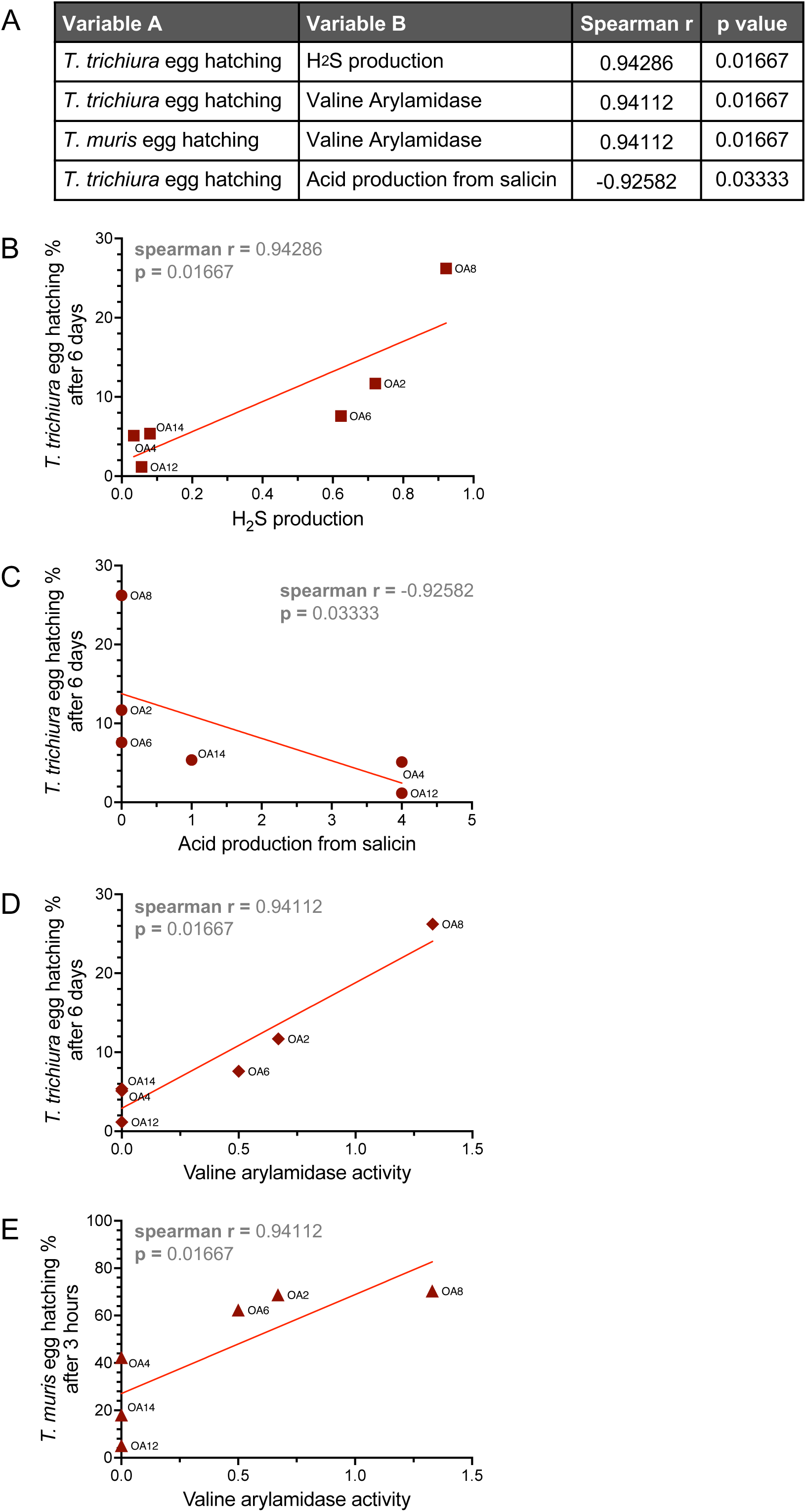
H_2_S, salicin acidification, and valine arylamidase are associated with *Trichuris egg* hatching by bacteria. (A) Spearman r and p values corresponding to significant correlations between *Trichuris* egg hatching and other measured variables of the OA isolates. (B) Relationship between H_2_S production by the OA isolates and *T. trichiura* egg hatching rate. (C) Relationship between acid production from salicin by the OA isolates and *T. trichiura* egg hatching rate. (D) Relationship between valine arylamidase activity by the OA isolates and *T. trichiura* egg hatching rate. (E) Relationship between valine arylamidase activity by the OA isolates and *T. muris* egg hatching rate. Spearman r and p values are depicted on each plot. Each dot corresponds to the average value per OA isolate and the red line depicts the linear regression slope.

## Discussion

Our comprehensive enzymatic profiling of OA isolates coupled with analysis of SCFAs and H_2_S production revealed a broad range of metabolic activities that are associated with host physiology, including immunity. Consistent with this observation, several OA isolates increased the number and proportion of colonic iTregs when introduced into GF mice, comparable to levels achieved upon colonization with the mixture of 17 Clostridia strains used as our benchmark. SCFAs were previously associated with the ability of the Clostridia mixture to induce Tregs [5, 6]. We did not observe a correlation between SCFA production and iTreg induction by the OA isolates, indicating that SCFAs are not sufficient to distinguish Treg-inducing OA isolates. Subsequent to the initial characterization of the Clostridia mixture, other metabolites such as bile acids have been shown to mediate Treg differentiation in the presence of intestinal bacteria [81], which may contribute to Treg differentiation induced by OA isolates. Alternatively, it is possible that previously unknown factors are involved. For example, we identified strong positive correlations between Treg induction and alanine arylamidase and tyrosine arylamidase activities that warrant further study. Our observation that three different species, *R. hominis*-OA2, *P. sordellii*-OA8, and *E. hirae*-OA12, consistently exhibited the highest Treg and Th17 cell induction as well as arylamidase activity, suggests a predictive potential for these enzymes in identifying bacteria with immunomodulatory capacity. Overall, this is one of the few studies that have functionally profiled individual bacterial isolates from understudied populations beyond inferring properties from genomic data.

The ability of some OA isolates to induce both Tregs and Th17 cells is consistent with the co-regulation of these cell types. Thus, a model in which helminth-associated bacteria inhibit inflammatory diseases by shifting the balance towards immune tolerance may be too simplistic. Although our approach allowed us to examine properties of individual bacteria in isolation and reduce variables, it would be interesting to determine whether the presence of *Trichuris* alters the properties of OA isolates, or vice versa. Along these lines, we noted correlations between *Trichuris* egg hatching as determined in our prior study, and H_2_S production and valine arylamidase activity. In addition, the helminth-associated *Peptostreptococcaceae* isolates *R. hominis*-OA2, *R. hominis*-OA6, and *P. sordellii*-OA8 had high glutamic acid decarboxylase (GAD) activity which is lacking in *E. hirae*-OA12, an isolate that fails to induce *T. muris* and *T. trichiura* egg hatching (Figure 2A) [45]. GAD catalyzes the formation of the neurotransmitter gamma-aminobutyric acid (GABA), and bacterially produced GAD and GABA have been shown to act on *Caenorhabditis elegans*, a nematode related to *Trichuris* [82]. *Trichuris* species are not genetically tractable. Therefore, we suggest that testing the role of bacterial products downstream of the above enzymatic pathways may lead to insights into the life cycle of this medically important parasite.

In conclusion, our findings describe the functional properties of bacteria isolated from helminth-colonized individuals in Malaysia and identify correlates of Treg induction and *Trichuris* egg hatching. Although our analyses focused on a finite set of parameters and bacterial taxa, they unveil key features of these specific helminth-associated bacteria that can be targeted for deeper analysis in future studies. As more bacteria from individuals in understudied populations become available for study along with their full genomes, it will be important to determine the extent to which these functional properties are represented in related taxa, which will enable us to infer how the selective presence of these bacteria in certain populations can impact their health status.

## Acknowledgements

We would like to thank Margie Alva, Juan Carrasquillo and David Basnight for their help in the NYU Gnotobiotic Facility, the NYU Flow Cytometry Core for training and access to equipment, and the NYU Reagent Preparation service for providing bacterial media. We would also like to thank Drew Jones, P’ng Loke, Victor Torres, Juan Lafaille, and members of the Cadwell and Loke Labs for their constructive comments. Figure 1 was created using BioRender.com. This work was in part funded by NIH grants DK093668 (K.C.), HL123340 (K.C.), AI130945 (K.C., P.L., Y.A.L.L.), AI140754 (K.C.), DK124336 (K.C.), AI121244 (K.C.), AI133977 (P.L.), 1DP2HD101401-01, (C.J.G.), DK135816-01 (C.J.G.), and 2T32AI007180 (S.S.). Further funding was provided by the Faculty Scholar grant from the Howard Hughes Medical Institute (K.C.), Crohn’s & Colitis Foundation (K.C.), Kenneth Rainin Foundation (K.C.), Judith & Stewart Colton Center of Autoimmunity (K.C.), SMRT Grant Competition award from Pacific Biosciences (K.C.), W.M. Keck Foundation (C.J.G.), Kenneth Rainin Foundation (C.J.G.), and RAPP funding from Weill Cornell Medicine (C.J.G.).

## Author Contributions

S.S. and K.C. conceived and designed the study. S.S. performed the experiments and analyzed and interpreted the data. A.L. helped perform enzyme assays and A.V. helped perform H_2_S assays. W.B.J. and C.J.G. performed and analyzed LC-MS to quantify SCFAs. D.E. performed the effect size analysis and assisted with computational and statistical analyses. K.C., P.L. and Y.A.L.L. oversaw interpretation of the data. S.S. and K.C. wrote the paper with input from all authors.

## Supplemental figures - titles and legends

**Figure S1.**
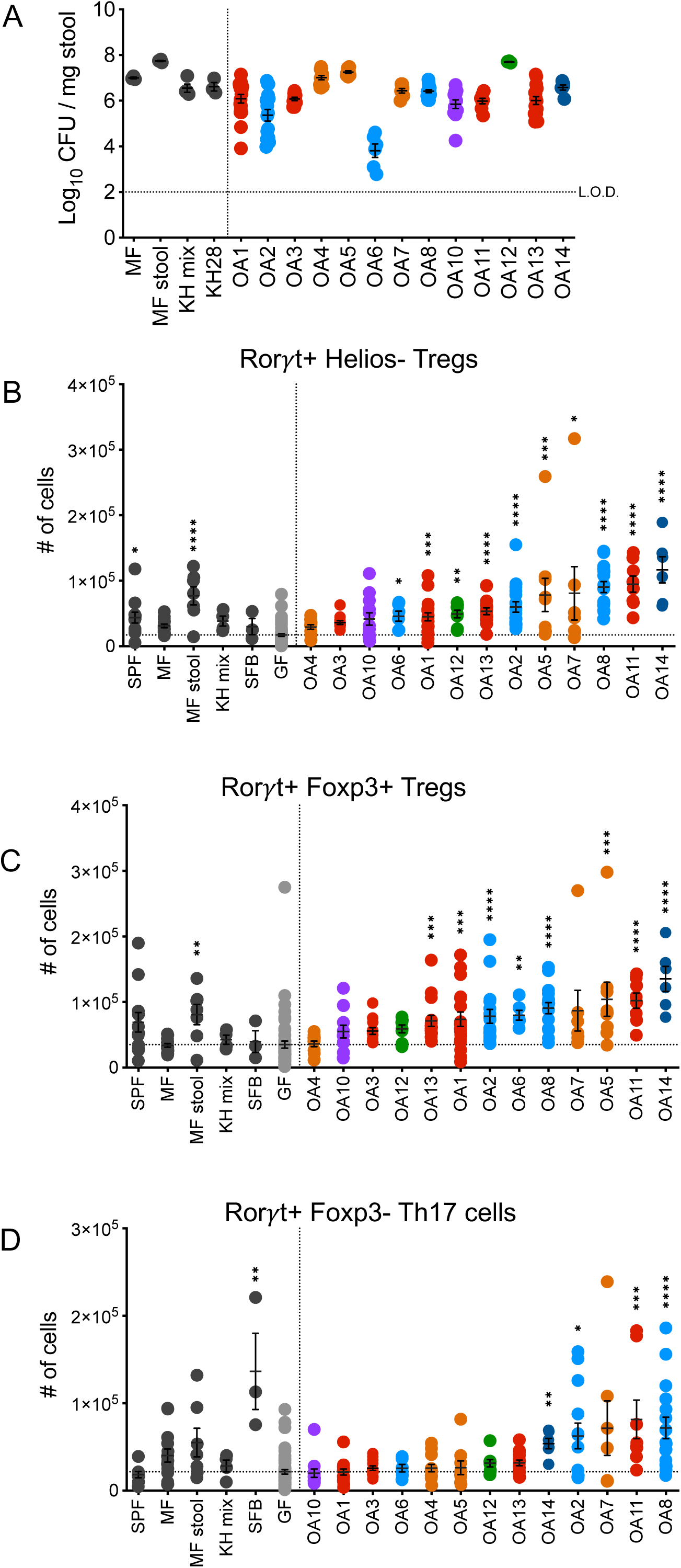
Bacterial burden and absolute numbers of Treg and Th17 cells induced by the OA isolates. (A) Colony forming units (CFUs) per mg of stool from mice colonized with indicated bacteria on the day cells were collected for analysis by flow cytometry. Each dot corresponds to one mouse. (B-D) Absolute numbers of Rorγt^+^ Helios^-^ Foxp3^+^ Tregs (B), Rorγt^+^ Foxp3^+^ Tregs (C) and Rorγt^+^ Foxp3^-^ Th17 cells (D) in the colon lamina propria of mice colonized with the indicated bacteria. SPF, Specific pathogen free; MF, Minimal flora mice; MF stool, GF mice gavaged with stool from Minimal flora mice; KH mix, Mice gavaged with consortium of 17 Clostridia strains from Atarashi et al., 2013; SFB, Segmented filamentous bacteria; GF, Germ-free. Each dot corresponds to one mouse. Each group was tested in at least 2 independent experiments consisting of 2-5 mice. Kruskal-Wallis with Dunn’s multiple comparisons test was used to compare between each group and GF in B-D. *p < 0.05, **p < 0.01, ***p < 0.001, ****p < 0.0001.

**Figure S2.**
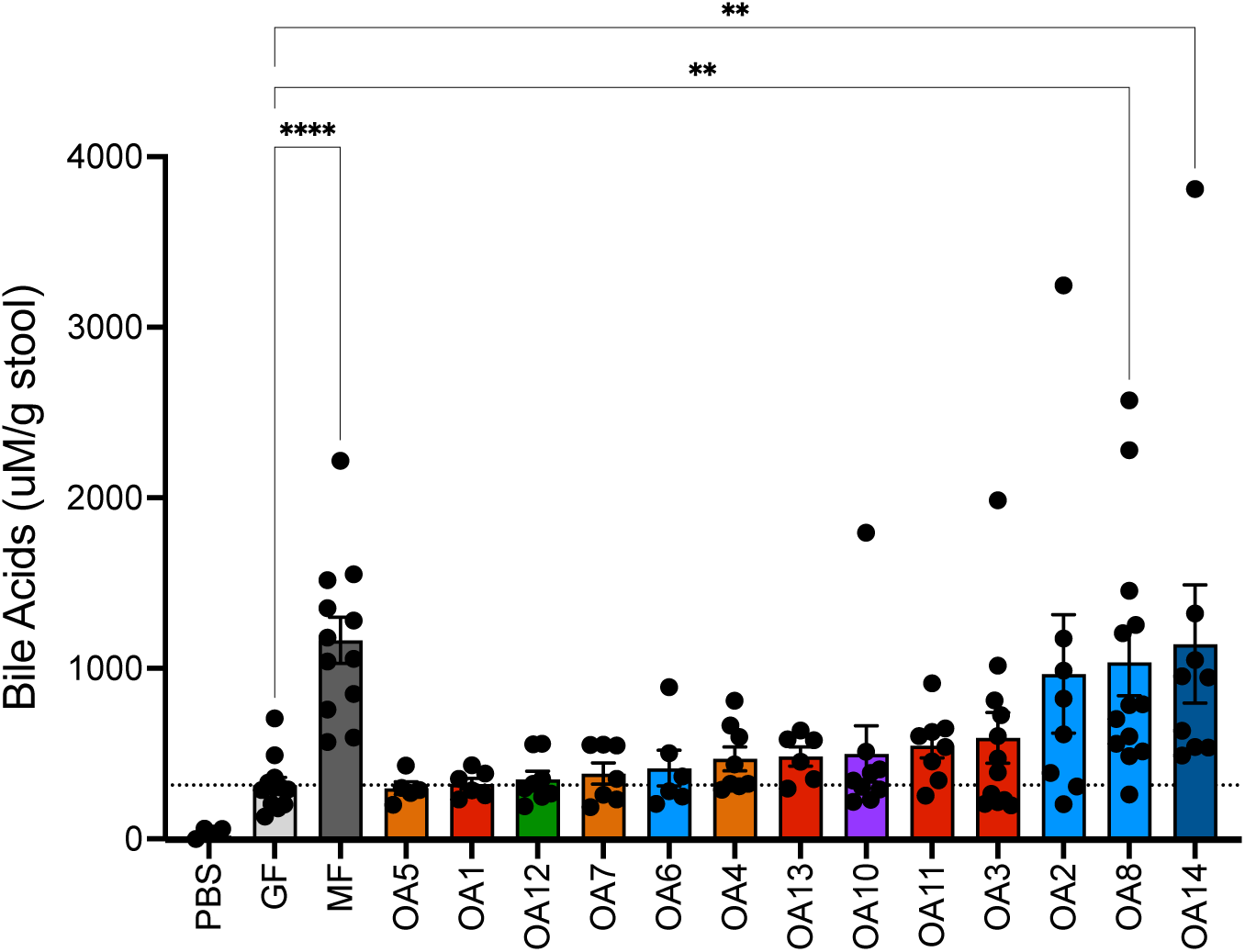
Monocolonization with a subset of OA isolates increases total bile acids in stool. Quantification of total bile acids per g of stool from mice monocolonized with the indicated bacteria. PBS, PBS as a technical control; GF, germ-free mouse; MF, minimal flora mouse. Kruskal-Wallis with Dunn’s multiple comparisons test was used to compare between each group and GF. **p < 0.01, ****p < 0.0001.

**Figure S3.**
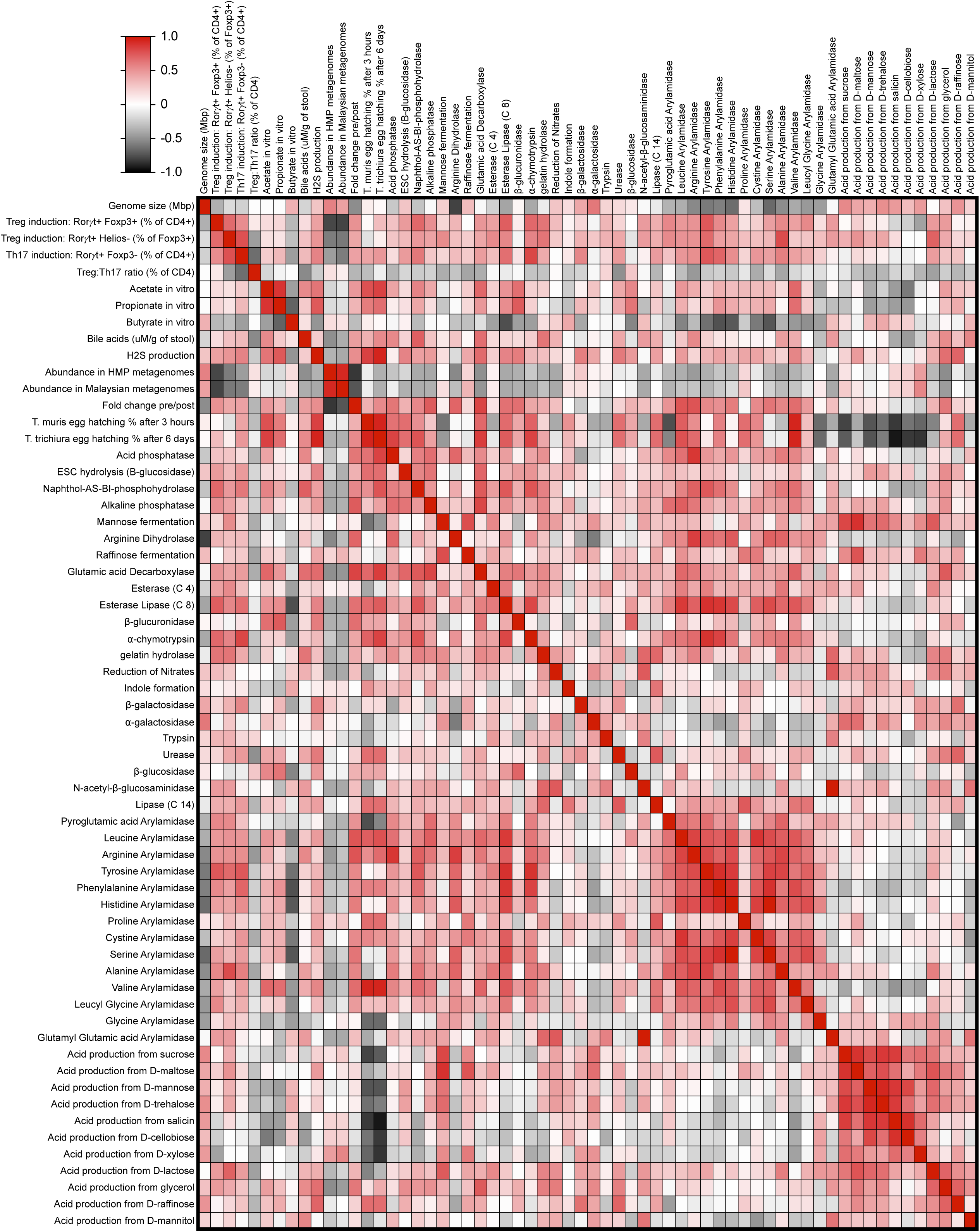
Spearman correlation matrix depicting relationships between OA isolate properties. Heatmap showing Spearman correlations between all measured variables for the OA isolates including Treg and Th17 cell induction, *Trichuris* egg hatching, and production of SCFAs, BAs, H_2_S, and enzymes. Variables with insufficient data points to complete the correlation analysis were removed from the heatmap.

**Figure S4.**
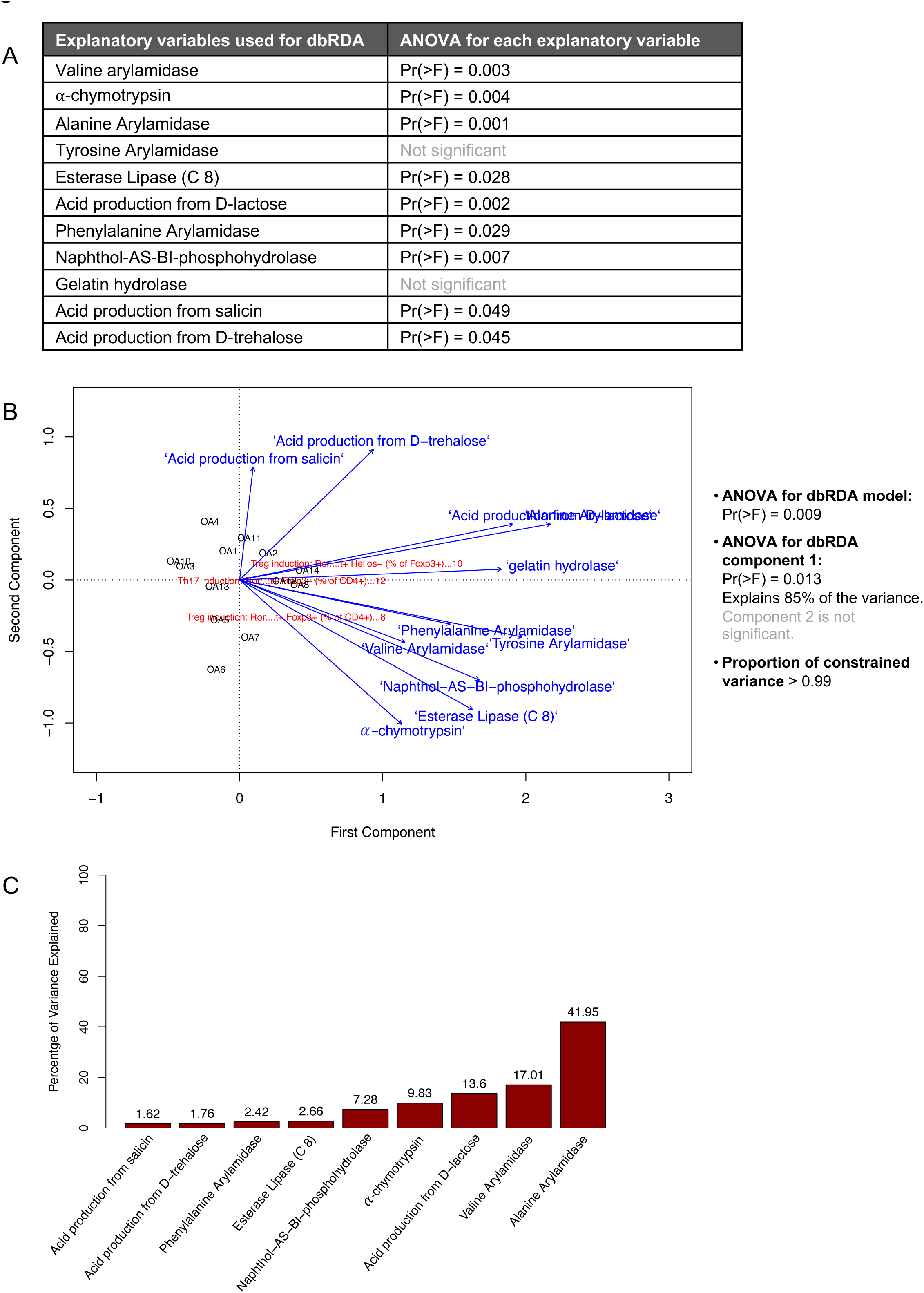
dbRDA analysis to determine effect size of bacterial enzyme production on lymphocyte differentiation induced by the OA isolates. (A) Enzyme production data used to perform the distance-based redundancy analysis (dbRDA) and statistical significance of the F statistic for each variable. (B) PCA biplot depicting dbRDA results for Treg and Th17 cell induction and enzyme production. (C) Proportion of variance explained by 10 significant variables on the variance of Treg and Th17 cell induction.

## Materials and Methods

### Gnotobiotic mice

Germfree (GF) C57BL/6J were bred and maintained in flexible-film isolators at the New York University Grossman School of Medicine Gnotobiotics Animal Facility. Absence of fecal bacteria was confirmed monthly by evaluating the presence of 16S DNA in stool samples by qPCR as previously described [83]. Minimal flora (MF) mice harboring the consortium of 15 bacteria described in [52] were kept in a separate isolator. For inoculation with bacteria, GF mice were housed in Bioexclusion cages (Tecniplast) with access to sterile food and water. An equal amount of male and female mice 6-8 weeks of age was used for all experiments. All animal studies were performed according to protocols approved by the NYU Grossman School of Medicine Institutional Animal Care and Use Committee.

### Bacterial strains

OA isolates were previously described [45]. The consortium of 17 Clostridia isolates (KH mix) including *Clostridium aldenense* (KH28) was kindly provided by K. Honda (RIKEN Center for Integrative Medical Sciences, Japan) [5]. All bacteria were cultured under anaerobic conditions in an anaerobic chamber (Coy Labs). Frozen glycerol stocks (30% glycerol) of all bacteria were prepared. Glycerol stocks of the OA isolates were streaked onto BRU agar plates (Anaerobe Systems) and incubated anaerobically for 48 hours at 37°C. PYG broth (Anaerobe Systems) inoculated with single colonies was grown at 37°C. OA1, 2, 4, 5, 6, 8, 11, 12, 13, 14, the KH mix, and KH28 were grown for 24 hours. OA3, 7, 9, and 10 required 3 days to reach similar turbidity. To quantify colony forming units, we performed serial dilutions of liquid culture in sterile PBS and plated on BRU agar plates. Segmented Filamentous Bacteria (SFB) were kindly provided by D. Littmann (NYU Grossman School of Medicine) [69] in the form of stool from GF mice monocolonized with SFB. SFB burden was confirmed in stool by qPCR as described in [69].

### Enzyme activity assays

OA isolates were grown anaerobically for 24 hours or 3 days, prepared as per manufacturer’s instructions, and inoculated onto test strips from APIⓇ ZYM, APIⓇ Rapid ID 32A, and APIⓇ 20A Microbial Identification Kits (BioMerieux). Test strips were incubated at 37°C for either 4 hours or 24 hours and at either aerobic or anaerobic conditions depending on the manufacturer’s instructions, after which enzyme reactions were assessed visually based on colorimetric changes.

### Quantification of SCFAs using LC-MS

Bacterial cultures of OA isolates were spun down at 14,000g for 5 minutes at 4°C and supernatant was collected and stored at -80°C. Cecal contents were harvested from GF mice, MF mice, or mice monocolonized with OA2 and stored at -80°C. For the measurement of SCFAs in bacterial liquid culture, a 10 µL aliquot of the culture was mixed with 190 µL of short-chain fatty acids (SCFAs) derivatization solution (1 mM 2,2′-dipyridyl disulfide, 1 mM triphenylphosphine, and 1 mM 2-hydrazinoquinoline dissolved in acetonitrile). For the measurement of SCFAs in cecal contents, ∼10mg of cecal sample was resuspended in 50 µL of 50% MeOH (in H2O) and vortexed for 10 min (some beads were added to disperse the cecal material). Then the mixture was spun down, and 10 µL of supernatant was mixed with 190 µL SCFAs derivatization solution.

For both bacterial liquid culture and cecal content samples, the resulting mixtures were vortexed and incubated at 60 °C for 1 hr. The mixture was centrifuged at 21000 x g for 20 min, and the supernatant was analyzed using an Agilent 1290 LC system coupled to an Agilent 6530 quadrupole time-of-flight (QTOF) mass spectrometer with a 130Å, 1.7 μm, 2.1 mm × 100 mm ACQUITY UPLC BEH C18 column (Waters). We used the following solvent system: A: H2O with 0.1% formic acid; B: Methanol with 0.1% formic acid. 1 µL of each sample was injected, and the flow rate was 0.35 mL/min with a column temperature of 40 °C. The gradient for HPLC-MS analysis was: 0-6.0 min, 99.5%-70.0% A; 6.0-9.0 min, 70.0%-2.0% A; 9.0-9.4 min, 2.0% A; 9.4-9.6min, 2.0%-99.5% A. Peaks were assigned by comparison with authentic standards.

### Hydrogen sulfide production assay

In a 24-well plate, 10ul of 24-hour or 3-day bacterial culture was added to 1ml of sterile PYG media. Each plate was covered with a piece of lead acetate paper cut to fit the 24-well plate, followed by the plate lid over the lead acetate paper. After 24 hours, photos were taken to quantify darkening of the lead acetate paper using Image J.

### Quantification of Treg and Th17 cell induction

Male and female germ-free C57BL/6J mice were monocolonized at 6-8 weeks of age by oral gavage with ∼1×10^7^ colony forming units (CFU) of indicated bacteria. 21-28 days later, mice were euthanized, and the colon and cecum were harvested. For single cell suspension, colonic and cecal tissues were flushed with HBSS (Gibco), fat and Peyer’s patches were removed, and the tissue was cut into 5-6 pieces. Tissue bits were incubated first with 20 mL of HBSS with 2% HEPES (Corning), 1% sodium pyruvate (Corning), 5mM EDTA, and 1 mM dithiothreitol (Sigma-Aldrich) for 15 min at 37°C with shaking, and then with new 20 mL of HBSS with 2% HEPES, 1% sodium pyruvate, 5mM EDTA for 10 min at 37°C with shaking. Tissue bits were washed in HBSS + 5% FCS, minced, and then enzymatically digested with collagenase D (0.5 mg/mL, Roche) and DNase I (0.01 mg/mL, Sigma-Aldrich) for 30-45 min at 37°C with shaking. Digested solutions were passed through a 70 mm cell strainer (BD) and cells were subjected to gradient centrifugation using 40% Percoll (Sigma-Aldrich).

Surface and transcription factor staining was performed per manufacturer’s instructions in PBS + 2% FBS for 20 min on ice. Zombie Aqua Fixable Viability Kit (Biolegend) was used to exclude dead cells. Surface markers were stained with anti-CD45 Pacific Blue, anti-CD3 FITC and anti-CD4 APC-Cy7 from Biolegend. For intracellular staining of transcription factors, cells were permeabilized with the eBioscience Foxp3/Transcription Factor Staining Buffer Set (Thermo Fisher Scientific) at room temperature for 30 min, and then stained with anti-Foxp3 APC and anti-Rorγt PE from Invitrogen, and anti-Helios PE-Cy7 from Biolegend. Samples were acquired on the CytoFLEX analyzer (Beckman Coulter) and analyzed using FlowJo 10.8.1.

### Quantification of bile acids in stool samples

10-15mg of stool was homogenized in cold 100% ethanol at a final concentration of 10mg/ml using Zirconia/Silica 1.0mm beads. Homogenates were spun down at 21,000g for 3 minutes at 4°C and 450ul of the supernatant was transferred to a clean 1.5ml tube. The sample was then dried down in a speedvac before reconstituting in ultrapure water. Bile acid concentration in each sample was quantified using the Bile Acid Assay Kit (Sigma #MAK309) following the manufacturer’s instructions, and the SpectraMax M3 Multi-Mode Microplate Reader with SoftMax Pro 6.5.1. software.

### Distance-based redundancy analysis (dbRDA)

Enzyme production levels were normalized, and one enzyme (⍺-mannosidase) that had zero production for all isolates was removed, for further statistical analyses. Least absolute shrinkage and selection operator (LASSO) regression was used as a feature selection method of enzymatic production. Enzymes that contributed significantly to prediction of Treg and Th17 induction and enzymes that had significant Spearman correlation coefficients were chosen as explanatory variables for dbRDA. Both analyses were conducted using R version 4.2.2.

### Statistical analysis

For *in vitro* and *in vivo* experiments, the number of repeats per group is annotated in corresponding figure legends. Significance for all experiments was assessed using GraphPad Prism software. Specific tests are annotated in corresponding figure legends. P values correlate with symbols: ns or no symbol = not significant, *p < 0.05, **p < 0.01, ***p < 0.001, ****p < 0.0001.

---

**Table S1.** Least absolute shrinkage and selection operator (LASSO) regression was performed to identify variables that could predict Treg and Th17 cell induction by the OA isolates. R^2^ values for the model are depicted next to each cell population type, and important variables and their corresponding coefficients are below.

| **Treg induction: Roryt+ Foxp3+ (% of CD4+), $R^2 > 0.3$** |  |
| --- | --- |
| <u>Variable</u> | <u>Coefficient</u> |
| Esterase Lipase (C 8) | 1.73E-02 |
| **Treg induction: Roryt+ Helios- (% of Foxp3+), $R^2 > 0.99$** |  |
| <u>Variable</u> | <u>Coefficient</u> |
| Alanine Arylamidase | 3.44E-01 |
| Alpha-Chymotrypsin | 6.94E-02 |
| Beta-galactosidase | 4.02E-02 |
| Alpha-Arabinosidase | -3.88E-02 |
| Gelatin Hydrolase | 3.27E-02 |
| Acid production from L-rhamnose | 2.97E-02 |
| Acid production from D-trehalose | 2.92E-02 |
| Acid production from D-maltose | 2.68E-02 |
| Acid production from D-glucose | 1.64E-02 |
| Pyroglutamic Acid Arylamidase | 1.33E-02 |
| **Th17 cell induction: Roryt+ Foxp3- (% of CD4+), $R^2 > 0.97$** |  |
| <u>Variable</u> | <u>Coefficient</u> |
| Beta-Galactosidase | 1.24E-01 |
| Beta-Galactosidase 6 Phosphate | -5.88E-02 |
| Arginine Arylamidase | 4.77E-02 |
| Alpha-Chymotrypsin | 3.19E-02 |
| Alanine Arylamidase | 2.19E-02 |
| Cysteine Arylamidase | 1.12E-02 |
| Acid Production from D-trehalose | 1.10E-02 |
| Acid Production from D-sorbitol | 4.18E-03 |
| ESC Hydrolysis (B-glucosidase) | 3.94E-04 |

## References

1. Turnbaugh, P.J., et al., The human microbiome project. Nature, 2007. 449(7164): p. 804–10.

2. Donaldson, G.P., S.M. Lee, and S.K. Mazmanian, Gut biogeography of the bacterial microbiota. Nat Rev Microbiol, 2016. 14(1): p. 20–32.

3. Macpherson, A.J. and N.L. Harris, Interactions between commensal intestinal bacteria and the immune system. Nat Rev Immunol, 2004. 4(6): p. 478–85.

4. Ruff, W.E., T.M. Greiling, and M.A. Kriegel, Host–microbiota interactions in immune-mediated diseases. Nature Reviews Microbiology, 2020. 18(9): p. 521–538.

5. Atarashi, K., et al., Treg induction by a rationally selected mixture of Clostridia strains from the human microbiota. Nature, 2013. 500(7461): p. 232–6.

6. Narushima, S., et al., Characterization of the 17 strains of regulatory T cell-inducing human-derived Clostridia. Gut Microbes, 2014. 5(3): p. 333–9.

7. Belkaid, Y. and T.W. Hand, Role of the microbiota in immunity and inflammation. Cell, 2014. 157(1): p. 121–41.

8. Ivanov, II and K. Honda, Intestinal commensal microbes as immune modulators. Cell Host Microbe, 2012. 12(4): p. 496–508.

9. Sun, M., et al., Microbiota metabolite short chain fatty acids, GPCR, and inflammatory bowel diseases. J Gastroenterol, 2017. 52(1): p. 1–8.

10. Clemente, J.C., et al., The microbiome of uncontacted Amerindians. Sci Adv, 2015. 1(3).

11. Fragiadakis, G.K., et al., Links between environment, diet, and the hunter-gatherer microbiome. Gut Microbes, 2019. 10(2): p. 216–227.

12. Groussin, M., et al., Elevated rates of horizontal gene transfer in the industrialized human microbiome. Cell, 2021. 184(8): p. 2053–2067.e18.

13. Sonnenburg, E.D. and J.L. Sonnenburg, The ancestral and industrialized gut microbiota and implications for human health. Nat Rev Microbiol, 2019. 17(6): p. 383–390.

14. McCall, L.I., et al., Home chemical and microbial transitions across urbanization. Nat Microbiol, 2020. 5(1): p. 108–115.

15. Vangay, P., et al., US Immigration Westernizes the Human Gut Microbiome. Cell, 2018. 175(4): p. 962–972.e10.

16. De Filippo, C., et al., Impact of diet in shaping gut microbiota revealed by a comparative study in children from Europe and rural Africa. Proc Natl Acad Sci U S A, 2010. 107(33): p. 14691–6.

17. Yatsunenko, T., et al., Human gut microbiome viewed across age and geography. Nature, 2012. 486(7402): p. 222-7.

18. Jha, A.R., et al., Gut microbiome transition across a lifestyle gradient in Himalaya. PLOS Biology, 2018. 16(11): p. e2005396.

19. Abdill, R.J., E.M. Adamowicz, and R. Blekhman, Public human microbiome data are dominated by highly developed countries. PLOS Biology, 2022. 20(2): p. e3001536.

20. Pasolli, E., et al., Extensive Unexplored Human Microbiome Diversity Revealed by Over 150,000 Genomes from Metagenomes Spanning Age, Geography, and Lifestyle. Cell, 2019. 176(3): p. 649–662.e20.

21. Kupritz, J., et al., Helminth-Induced Human Gastrointestinal Dysbiosis: a Systematic Review and Meta-Analysis Reveals Insights into Altered Taxon Diversity and Microbial Gradient Collapse. mBio, 2021. 12(6): p. e0289021.

22. Rapin, A. and N.L. Harris, Helminth-Bacterial Interactions: Cause and Consequence. Trends Immunol, 2018. 39(9): p. 724–733.

23. Loke, P.n. and N.L. Harris, Networking between helminths, microbes, and mammals. Cell Host & Microbe, 2023. 31(4): p. 464–471.

24. Zaiss, M.M. and N.L. Harris, Interactions between the intestinal microbiome and helminth parasites. Parasite Immunol, 2016. 38(1): p. 5–11.

25. Walusimbi, B., et al., The effects of helminth infections on the human gut microbiome: a systematic review and meta-analysis. Frontiers in Microbiomes, 2023. 2.

26. Bethony, J., et al., Soil-transmitted helminth infections: ascariasis, trichuriasis, and hookworm. Lancet, 2006. 367(9521): p. 1521–32.

27. WHO, Soil-Transmitted Helminth Infections. WHO Fact Sheets. 2020, World Health Organization: Geneva, Switzerland.

28. Jourdan, P.M., et al., Soil-transmitted helminth infections. Lancet, 2018. 391(10117): p. 252–265.

29. Weinstock, J.V. and D.E. Elliott, Helminths and the IBD hygiene hypothesis. Inflamm Bowel Dis, 2009. 15(1): p. 128–33.

30. Bach, J.F., The hygiene hypothesis in autoimmunity: the role of pathogens and commensals. Nat Rev Immunol, 2017.

31. Elliott, D.E. and J.V. Weinstock, Helminth-host immunological interactions: prevention and control of immune-mediated diseases. Ann N Y Acad Sci, 2012. 1247: p. 83–96.

32. Finlay, C.M., K.P. Walsh, and K.H. Mills, Induction of regulatory cells by helminth parasites: exploitation for the treatment of inflammatory diseases. Immunol Rev, 2014. 259(1): p. 206–30.

33. McSorley, H.J. and R.M. Maizels, Helminth infections and host immune regulation. Clin Microbiol Rev, 2012. 25(4): p. 585–608.

34. Hang, L., et al., Heligmosomoides polygyrus bakeri infection activates colonic Foxp3+ T cells enhancing their capacity to prevent colitis. J Immunol, 2013. 191(4): p. 1927–34.

35. D’Elia, R., et al., Regulatory T cells: a role in the control of helminth-driven intestinal pathology and worm survival. J Immunol, 2009. 182(4): p. 2340–8.

36. Klementowicz, J.E., M.A. Travis, and R.K. Grencis, Trichuris muris: a model of gastrointestinal parasite infection. Semin Immunopathol, 2012. 34(6): p. 815–28.

37. Maizels, R.M., Regulation of immunity and allergy by helminth parasites. Allergy, 2020. 75(3): p. 524–534.

38. Maizels, R.M., H.H. Smits, and H.J. McSorley, Modulation of Host Immunity by Helminths: The Expanding Repertoire of Parasite Effector Molecules. Immunity, 2018. 49(5): p. 801–818.

39. Loke, P. and Y.A. Lim, Helminths and the microbiota: parts of the hygiene hypothesis. Parasite Immunol, 2015. 37(6): p. 314–23.

40. Lawson, M.A.E., I.S. Roberts, and R.K. Grencis, The interplay between Trichuris and the microbiota. Parasitology, 2021: p. 1–8.

41. Zaiss, M.M., et al., The Intestinal Microbiota Contributes to the Ability of Helminths to Modulate Allergic Inflammation. Immunity, 2015. 43(5): p. 998–1010.

42. Ramanan, D., et al., Helminth infection promotes colonization resistance via type 2 immunity. Science, 2016. 352(6285): p. 608–12.

43. Lopetuso, L.R., et al., Commensal Clostridia: leading players in the maintenance of gut homeostasis. Gut Pathogens, 2013. 5(1): p. 23.

44. Atarashi, K., et al., Induction of colonic regulatory T cells by indigenous Clostridium species. Science, 2011. 331(6015): p. 337–41.

45. Sargsian, S., et al., Clostridia isolated from helminth-colonized humans promote the life cycle of Trichuris species. Cell Rep, 2022. 41(9): p. 111725.

46. Maria, R., et al., Evaluation of Antibacterial Properties of Organic Gutta-percha Solvents and Synthetic Solvents Against Enterococcus faecalis. J Int Soc Prev Community Dent, 2021. 11(2): p. 179–183.

47. Suchomel, M., et al., Enterococcus hirae, Enterococcus faecium and Enterococcus faecalis show different sensitivities to typical biocidal agents used for disinfection. J Hosp Infect, 2019. 103(4): p. 435–440.

48. Koh, A., et al., From Dietary Fiber to Host Physiology: Short-Chain Fatty Acids as Key Bacterial Metabolites. Cell, 2016. 165(6): p. 1332–1345.

49. Arpaia, N., et al., Metabolites produced by commensal bacteria promote peripheral regulatory T-cell generation. Nature, 2013. 504(7480): p. 451–5.

50. Furusawa, Y., et al., Commensal microbe-derived butyrate induces the differentiation of colonic regulatory T cells. Nature, 2013. 504(7480): p. 446–50.

51. Smith, P.M., et al., The microbial metabolites, short-chain fatty acids, regulate colonic Treg cell homeostasis. Science, 2013. 341(6145): p. 569–73.

52. Brugiroux, S., et al., Genome-guided design of a defined mouse microbiota that confers colonization resistance against Salmonella enterica serovar Typhimurium. Nat Microbiol, 2016. 2: p. 16215.

53. Dallari, S., et al., Enteric viruses evoke broad host immune responses resembling those elicited by the bacterial microbiome. Cell Host Microbe, 2021. 29(6): p. 1014–1029.e8.

54. Cui, C., et al., CD4(+) T-Cell Endogenous Cystathionine γ Lyase-Hydrogen Sulfide Attenuates Hypertension by Sulfhydrating Liver Kinase B1 to Promote T Regulatory Cell Differentiation and Proliferation. Circulation, 2020. 142(18): p. 1752–1769.

55. Yang, R., et al., Hydrogen Sulfide Promotes Tet1-and Tet2-Mediated Foxp3 Demethylation to Drive Regulatory T Cell Differentiation and Maintain Immune Homeostasis. Immunity, 2015. 43(2): p. 251–63.

56. Braccia, D.J., et al., The Capacity to Produce Hydrogen Sulfide (H2S) via Cysteine Degradation Is Ubiquitous in the Human Gut Microbiome. Frontiers in Microbiology, 2021. 12.

57. Fiorucci, S., et al., Inhibition of hydrogen sulfide generation contributes to gastric injury caused by anti-inflammatory nonsteroidal drugs. Gastroenterology, 2005. 129(4): p. 1210–24.

58. Rahman, M.A., et al., Hydrogen sulfide dysregulates the immune response by suppressing central carbon metabolism to promote tuberculosis. Proc Natl Acad Sci U S A, 2020. 117(12): p. 6663–6674.

59. Fiorucci, S., et al., Enhanced activity of a hydrogen sulphide-releasing derivative of mesalamine (ATB-429) in a mouse model of colitis. Br J Pharmacol, 2007. 150(8): p. 996–1002.

60. Tanoue, T., K. Atarashi, and K. Honda, Development and maintenance of intestinal regulatory T cells. Nat Rev Immunol, 2016. 16(5): p. 295–309.

61. Sefik, E., et al., Individual intestinal symbionts induce a distinct population of RORgamma(+) regulatory T cells. Science, 2015. 349(6251): p. 993–7.

62. Ohnmacht, C., et al., The microbiota regulates type 2 immunity through RORgt+ T cells. Science, 2015. 349(6251): p. 989–993.

63. Yang, B.H., et al., Foxp3(+) T cells expressing RORgammat represent a stable regulatory T-cell effector lineage with enhanced suppressive capacity during intestinal inflammation. Mucosal Immunol, 2016. 9(2): p. 444–57.

64. Kanamori, M., et al., Induced Regulatory T Cells: Their Development, Stability, and Applications. Trends Immunol, 2016. 37(11): p. 803–811.

65. Bilate, A.M. and J.J. Lafaille, Induced CD4+Foxp3+ regulatory T cells in immune tolerance. Annu Rev Immunol, 2012. 30: p. 733–58.

66. Geva-Zatorsky, N., et al., Mining the Human Gut Microbiota for Immunomodulatory Organisms. Cell, 2017. 168(5): p. 928–943 e11.

67. Lathrop, S.K., et al., Peripheral education of the immune system by colonic commensal microbiota. Nature, 2011. 478(7368): p. 250–4.

68. Round, J.L. and S.K. Mazmanian, Inducible Foxp3+ regulatory T-cell development by a commensal bacterium of the intestinal microbiota. Proc Natl Acad Sci U S A, 2010. 107(27): p. 12204–9.

69. Ivanov, II, et al., Induction of intestinal Th17 cells by segmented filamentous bacteria. Cell, 2009. 139(3): p. 485–98.

70. Hirahara, K. and T. Nakayama, CD4+ T-cell subsets in inflammatory diseases: beyond the Th1/Th2 paradigm. International immunology, 2016. 28(4): p. 163–171.

71. Honda, K. and D.R. Littman, The microbiota in adaptive immune homeostasis and disease. Nature, 2016. 535(7610): p. 75–84.

72. Hang, S., et al., Bile acid metabolites control TH17 and Treg cell differentiation. Nature, 2019. 576(7785): p. 143–148.

73. Li, W., et al., A bacterial bile acid metabolite modulates T(reg) activity through the nuclear hormone receptor NR4A1. Cell Host Microbe, 2021. 29(9): p. 1366–1377.e9.

74. Paik, D., et al., Human gut bacteria produce ΤΗ17-modulating bile acid metabolites. Nature, 2022. 603(7903): p. 907–912.

75. Song, X., et al., Microbial bile acid metabolites modulate gut RORγ+ regulatory T cell homeostasis. Nature, 2020. 577(7790): p. 410–415.

76. Zhou, D., et al., Chymotrypsin both directly modulates bacterial growth and asserts ampicillin degradation-mediated protective effect on bacteria. Annals of Microbiology, 2013. 63(2): p. 623–631.

77. Miyoshi, S.-I., Extracellular proteolytic enzymes produced by human pathogenic vibrio species. Frontiers in Microbiology, 2013. 4.

78. Hurley, L.C. and R.I. Sommerville, Reversible inhibition of hatching of infective eggs of Ascaris suum (Nematoda). International Journal for Parasitology, 1982. 12(5): p. 463–465.

79. Sonowal, R., et al., Hydrolysis of aromatic β-glucosides by non-pathogenic bacteria confers a chemical weapon against predators. Proceedings. Biological sciences, 2013. 280(1762): p. 20130721–20130721.

80. Dziekońska-Rynko, J. and J. Rokicki, Activity of selected hydrolases in excretion-secretion products and extracts of adult Contracaecum rudolphii. Wiad Parazytol, 2005. 51(3): p. 227–31.

81. Larabi, A.B., H.L.P. Masson, and A.J. Bäumler, Bile acids as modulators of gut microbiota composition and function. Gut Microbes, 2023. 15(1): p. 2172671.

82. Urrutia, A., et al., Bacterially produced metabolites protect C. elegans neurons from degeneration. PLoS Biol, 2020. 18(3): p. e3000638.

83. Kernbauer, E., Y. Ding, and K. Cadwell, An enteric virus can replace the beneficial function of commensal bacteria. Nature, 2014. 516(7529): p. 94–8.

